# cvlr:Finding heterogeneously methylated genomic regions using ONT reads

**DOI:** 10.1101/2022.04.19.488395

**Authors:** Emanuele Raineri, Mariona Alberola i Pla, Marc Dabad, Simon Heath

## Abstract

**Summary:** Nanopore reads encode information on the methylation status of cytosines in `CpG` dinucleotides. The length of the reads makes it comparatively easy to look at patterns consisting of multiple loci; here we exploit this property to look for regions where one can define subpopulations of cells based on methylation patterns. As a benchmark we run our clustering algorithm on known imprinted genes and show that the clustering based on methylation is consistent with the phasing of the genome; we then scan chromosome 15 looking for windows corresponding to heterogeneous methylation. We can also compute the covariance of methylation across these regions while keeping into account the mixture of different types of reads.

**Availability:** https://github.com/EmanueleRaineri/releases

**Supplementary information:** Tables, figures, and some further explanations of the algorithms are available as online supplementary information.

## 1 Introduction

In this manuscript we exploit the fact that one can measure both DNA sequence and cytosine methylation status in one go via Nanopore sequencing [15] to scan the genome looking for regions where methylation changes between cells. The idea of quantifying the variability in binary epigenetic patterns in single reads was initially discussed in [10]. A probabilistic model to study allele specific methylation has been proposed by [7] on short reads measured with whole genome bisulfite sequencing; [6] discusses the analysis of RRBS libraries and [4] presents a statistical method for finding haplotpye specific methylation through long reads based on an Ising-like model but as far as we know there is no method for the analysis of methylation heterogeneity on long reads which does not rely on phasing, which is also used in [5] from where we take some benchmarking data. What we propose here is a software which can be run from the command line on the output of (among others) Nanopore sequencing experiments to cluster reads based on methylation patterns appearing on them. Since it does not rely on a previous phasing step, our method can capture allele specific differences but also other forms of heterogeneity. We show some results obtained by looking at chomosome 15, which we chose because it contains a number of well studied imprinted regions [13]. Our test data set is the sample `NA12878` sequenced by the nanopore consortium [9].

## 2 Results and Methods

To make sure we can recapitulate known allele specific methylation, we first looked at some genes known to be imprinted in many tissues (table 1). An example of analysis on GNAS is in figure 1. Supplementary material has more plots. For each gene we clustered in *k* = 2 groups the reads mapping on the corresponding genomic stretch, and also tagged each read with its haplotype as computed by `WhatsHap`[11]. We then check that each cluster is enriched in a specific haplotype. In some cases we do not look at the entire gene region, rather we selected stretches where the methylation seems to be at intermediate levels or positions surrounding known regulatory sites, or already described as imprinting control regions in [5]: the exact loci are under the “chr:start-end” heading.

**Figure 1:**
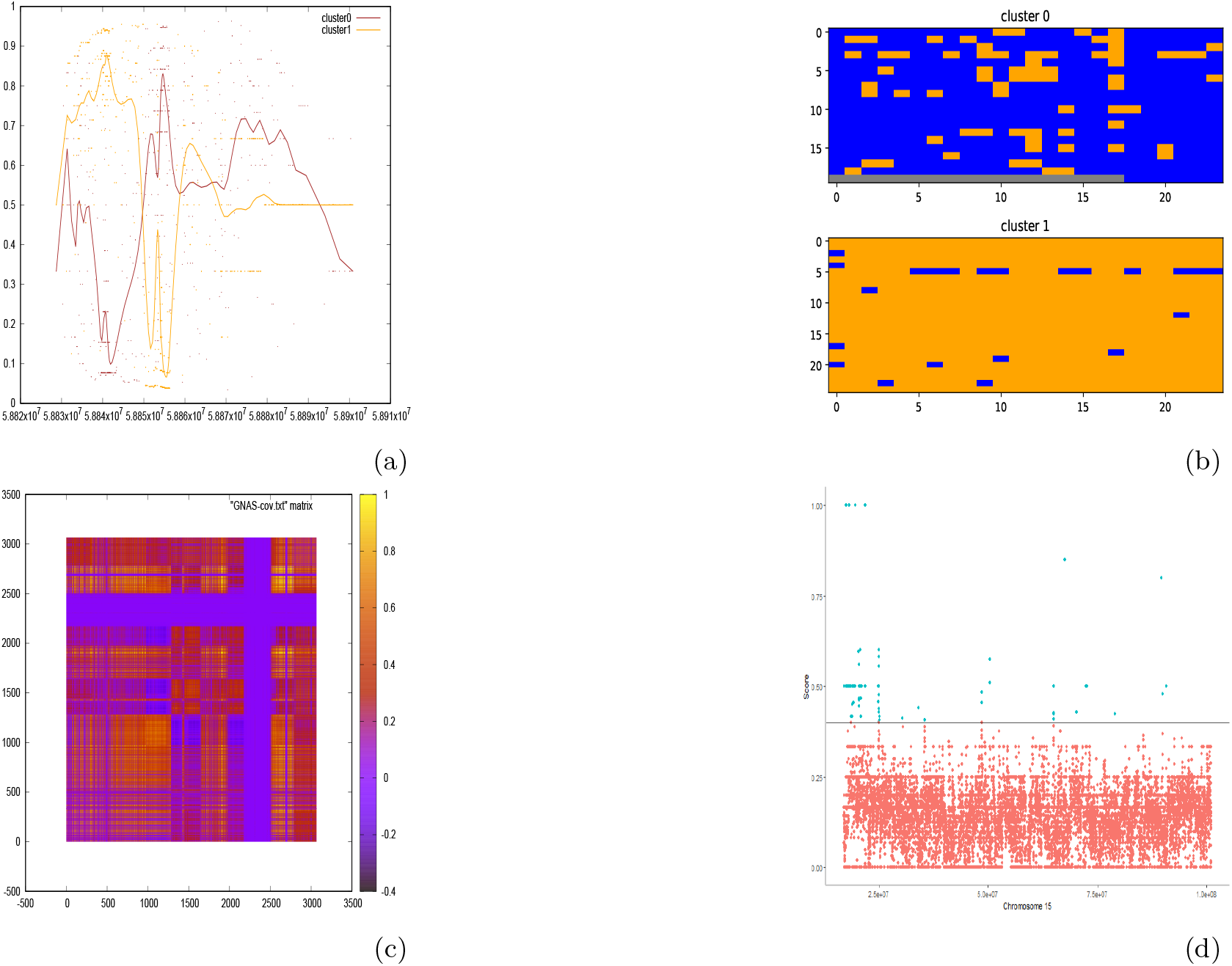
(a) Mean values of methylation in the 2 clusters over the GNAS region. (b) The two corresponding subpopulations of reads in the subregion 5.88545*e*7 – 5.88547*e*7. (blue = methylated, orange = not methylated, gray = not known). (c) Covariance matrix of the methylation in the GNAS gene. (d) Scan of chromosome 15 in windows of 5Kb. For each window we computed the clustering (*k* = 2) of the reads and the median absolute difference in methylation across clusters, and plotted it along the chromosome. Some of the peaks overlap with annotated genes and GM12878 enhancers.

**Table 1:** Known imprinted genes we looked at. For each region we repeat the clustering with different seeds for the random initialization and we pick the output with the best likelihood. The third column reports the median of the absolute difference in methylation between clusters; we compute its empirical p value (fourth column) using its distribution on a set of 1000 random cluster assignments. We also check that each cluster is enriched in a specific haplotype via a Fisher’s test (p value in the fifth column: a dash means that it was not possible to phase more than 20 reads mapping on the region.

| GENE | chr:start-end | $\Delta m$ | pval($\Delta m$) | pval (phasing) |
| --- | --- | --- | --- | --- |
| GNAS | chr20:58830000-58870000 | 0.614 | $< 10^{-3}$ | 1.6e-17 |
| H19dmr | chr11:1997582-2003510 | 0.175 | 0.023 | - |
| H19 | chr11:1995163-2001470 | 0.167 | 0.053 | - |
| PEG10 | chr7:94656325-94669695 | 0.233 | $< 10^{-3}$ | 2.1e-10 |
| MEG3 | chr14:100820000-100840000 | 0.167 | $< 10^{-3}$ | 0.044 |
| SNRPNdmr | chr15:24950000-24958000 | 0.497 | $< 10^{-3}$ | 3.3e-13 |
| IGF2dmr2 | chr11:2132761-2133882 | 0.111 | 0.03 | 0.13 |
| PLAGL1 | chr6:144000000-144020000 | 0.300 | $< 10^{-3}$ | 7e-05 |

We then divided chr15 in windows of 5Kb and clustered (with *k* = 2) the reads mapping on each window; The median absolute difference in methylation between the clusters at each locus is in figure 1d. We looked at 119 windows that have absolute difference greater than 0.4; 88 of them overlap with at least one of 33 genes and 1 enhancer. The software we used to produce figure 1 is called `cvlr` and it consists of three executables, `cvlr-cluster` which runs the clustering algorithm, `cvlr-meth-of-bam` which helps in extracting methylation information from `BAM` files and `cvlr-stats` which process the clusters to compute descriptive statistics. It is not necessary to use `cvlr-meth-of-bam` to produce the input for `cvlr-cluster`, users can equally adapt the output of eg `Nanopolish` [14]. Internally the algorithm sees the data as a binary matrix (with missing data) *X*, with *n* rows representing reads and *d* columns corresponding to genomic positions. Each position *X_ij_* can be 0 (unmethylated) 1 (methylated) or –1 (unknown). Reads are clustered (into *k* clusters) via a mixture of multivariate Bernoulli distributions. To this purpose we use an EM algorithm allowing for incomplete observations; the output of the algorithm is :

- a cluster assigment for each read
- a *k* × *d* matrix *μ* containing the average methylation for each cluster at each position
- the *d* × *d* covariance matrix of the methylation

The E step computes the posterior probability that a given read belongs to a certain cluster; the M step updates the *μ* matrix, replacing missing data with a weighted sum of the corresponding positions in the old *μ*. `cvlr-stats` computes, among other things, the median absolute difference in methylation and the posterior probability of difference in methylation between all pairs of clusters across all the positions (in pairs) using the method described in [12]. The command line allows to specify the number of clusters, a seed for the random initialization of the algorithm and the maximum number of iterations of the EM algorithm. We downloaded all the test data we used from [3] and processed them with our internal pipeline [2] which is based on `Megalodon` [1]. Enhancers were taken from [8]. In summary, we showed how to use `cvlr` to measure methylation heterogeneity along the human genome; although we used imprinted regions (and hence allele specific methylation) as a benchmark, `cvlr` can be run to detect subpopulation of reads regardless of whether they are due to an allelic effect and does not need a preliminary phasing step.

## 3 Acknowledgments

This work was funded by a Spanish Plan Nacional grant (project:PGC2018-099640-B-I00). Furthermore, we acknowledge support of the Spanish Ministry of Science and Innovation to the EMBL partnership, of the Centro de Excelencia Severo Ochoa and of the CERCA Programme / Generalitat de Catalunya. We also acknowledge support of the Spanish Ministry of Science and Innovation through the Instituto de Salud Carlos III of the Generalitat de Catalunya through Departament de Salut and Departament d’Universitats i Recerca. We also acknowledge the Co-financing with funds from the European Regional Development Fund by the Spanish Ministry of Science and Innovation corresponding to the Programa Operativo FEDER Plurirregional de Espana (POPE) 2014-2020 and by the Departament d’Universitats i Recerca of the Generalitat de Catalunya corresponding to the Programa Operatiu FEDER de Catalunya 2014-2020.

## Notes

https://github.com/EmanueleRaineri/releases

### Competing Interest Statement

The authors have declared no competing interest.

### Summary of Updates

We submitterd a wrong (old) version of the manuscript.

