## Supplementary Materials for "cvlr:Finding heterogeneously methylated genomic regions using ONT reads" (from another version of this preprint: version 1, 2022-04-19)

### 1 Analysis of known imprinted genes

## 1.1 H19

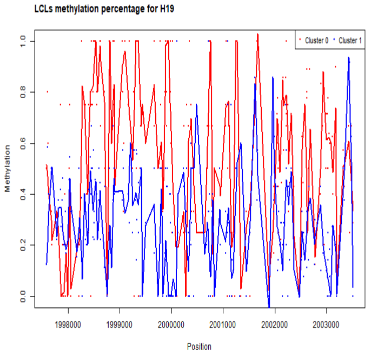


#### 1.2 IGF2

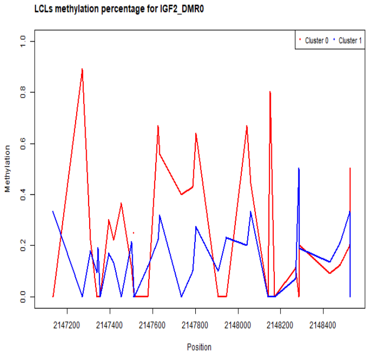


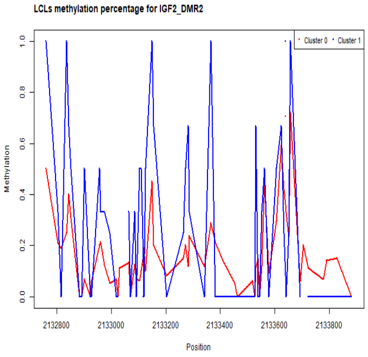

##### 1.3 MEG3

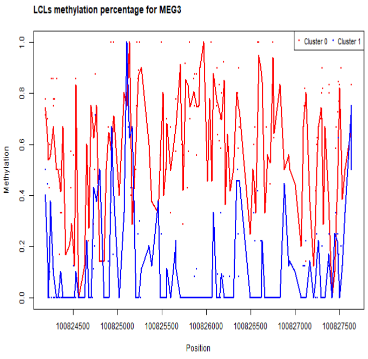


#### 2 Algorithms

##### 2.1 Mixture of multivariate bernoulli distribution

A mixture of multivariate Bernoulli distributions has the form

$$P(x_{ij}|\mu_{kj},\pi_k) = \prod_{i=1}^{N}\sum_{l=1}^{K}\pi_k\prod_{j=1}^{D}\mu_{kj}^{x_{ij}}(1-\mu_{kj})^{(1-x_{ij})}$$

where $\sum_k \pi_k = 1$ and $x_{ij} \in \{0,1\}$.

##### 2.2 likelihood

$$P(x|\theta) = \prod_{i=1}^{N} p(x_i|\mu) = \prod_{i=1}^{N}\sum_{l=1}^{K}\pi_l p(x_i|\mu_l) \tag{1}$$

Now, $x_i$ is a vector of dimension $D$, hence the above is equal to

$$\prod_{i=1}^{N}\sum_{l=1}^{K}\pi_l\prod_{j=1}^{D}p(x_{ij}|\mu_{lj})$$

Now take the logarithm

$$\sum_{i=1}^{N}\log\left(\sum_{l=1}^{K}\overbrace{\pi_l\prod_{j=1}^{D}p(x_{ij}|\mu_{lj})}^{\phi}\right) \tag{2}$$

Where I have introduced

$$\phi(\pi_l, x_i, \mu_l) = \pi_l \prod_{j=1}^{D} p(x_{ij}|\mu_{lj})$$

Hence I can rewrite (2) as

$$\sum_{i=1}^{N} \log \left( \sum_{l=1}^{K} \phi(\pi_l, x_i, \mu_l) \right)$$

which is equal to

$$\sum_{i=1}^{N} \log \left( \sum_{l=1}^{K} \exp \log \phi(\pi_l, x_i, \mu_l) \right) \tag{3}$$

If we introduce the `logsumexp` function

$$\text{lse}(s_1, \ldots, s_k) = \log(\exp s_1 + \ldots + \exp s_k) = \log(\sum_k \exp s_k)$$

we can rewrite (3) as

$$\sum_{i=1}^{N} \text{lse}(\log \phi(\pi_1, x_i, \mu_1), \ldots, \log \phi(\pi_K, x_i, \mu_K)) \tag{4}$$

### References
